# Distinct T cell chromatin landscapes in circulating lymphocytes from patients with systemic sclerosis

**DOI:** 10.1101/2021.01.10.426131

**Authors:** Diana R. Dou, Yang Zhao, Brian Abe, Rui Li, Lisa C. Zaba, Kathleen Aren, Mary Carns, Alec Peltekian, Natania Field, Lorinda S. Chung, Howard Y. Chang, Monique Hinchcliff

## Abstract

**Objectives:** Systemic sclerosis (SSc; scleroderma) disproportionately affects biological females, and results of multiple studies implicate lymphocyte derangements in disease pathogenesis supporting use of mycophenolate mofetil (MMF) treatment. Here, we surveyed chromatin accessibility of circulating CD4+ T lymphocytes from SSc patients commencing MMF to gain molecular insights into systemic immune activation.

**Methods:** Peripheral blood samples were collected longitudinally. We used the Assay for Transposase-Accessible Chromatin by sequencing (ATAC-seq) to interrogate genome-wide chromatin accessibility profiles of peripheral CD4+ T cells compared with publicly available healthy control (HC) data.

**Results:** ATAC-seq libraries were generated for 18 SSc patients [78% with diffuse cutaneous (dc), 78% female, and 17% + anticentromere autoantibodies (ACA)]. Disease status (SSc vs. HC), biological sex, and serum autoantibody type were significantly associated with CD4^+^ T cell epigenomic profile variability, while MMF treatment had no significant effect. Present serum ACAs associated with elevated T helper 2 (Th2) cell proportions. +ACA patients consistently displayed distinct epigenetic profiles of increased open chromatin at gene loci encoding fibrosis-driving Th2 cytokines IL-4, IL-13, and the IL-4 receptor.

**Conclusions:** Our results demonstrate the utility of interrogating chromatin accessibility profiles of patient CD4+ cells to stratify and understand better clinical SSc heterogeneity. They highlight a potential mechanism underlying the female sex preponderance for SSc as females had more open chromatin. Our findings nominate Th2 cell activation as a novel mechanistic hallmark and therapeutic opportunity to address SSc, especially in those with +ACA.

## INTRODUCTION

Emerging evidence regarding SSc pathogenesis suggests that vasculopathy, adaptive and innate autoimmunity, and fibrosis may develop concurrently (1). Adaptive immune involvement includes the presence of serum autoantibodies associated with SSc clinical manifestations [*e*.*g*., anticentromere antibodies (ACA) and pulmonary arterial hypertension (PAH)], and the presence of perivascular lymphocyte infiltrates, particularly in early skin disease. A recent finding that the epiregulin/EGFR+ fibroblast pathway promotes skin and lung fibrosis provides additional evidence of a role for innate immunity perturbations in SSc (2). Although mycophenolate mofetil (MMF) is commonly prescribed for SSc skin disease, with therapeutic effects attributed partly to modulating dermal macrophages (3), targeted therapies for skin disease in SSc remain elusive.

The majority of genomic profiling in SSc patients has focused on differential gene expression; however, chromatin accessibility is a critical regulator of gene expression and a key determinant of cellular identity. ATAC-seq enables sensitive interrogation of chromatin accessibility, particularly in rare hematopoietic populations (4-6). Additionally, ATAC-seq profiles in human immune cells more accurately reflect cell identity than transcriptomic profiles alone (6). Here, we performed bulk ATAC-seq on circulating CD4+ T cells from patients with active SSc skin disease, sampled longitudinally before and during MMF treatment. We evaluated associations between chromatin accessibility and clinical variables including sex, age, serum autoantibody profile, and MMF exposure.

## METHODS

### Clinical Cohort

The Northwestern University Institutional Review Board approved the study (IRB#STU00080199). All research participants provided informed consent in accordance with the Declaration of Helsinki. SSc patients fulfilled 2013 American College of Rheumatology SSc Classification Criteria (7) or ≥3 of 5 CREST features (calcinosis cutis, Raynaud phenomenon, esophageal dysmotility, sclerodactyly, telangiectasias). Blood samples were collected in sodium heparin tubes and shipped overnight at room temperature. A subset of patients initiated MMF for active skin disease at 250 mg twice daily with escalation to 1,000–1,500 mg twice daily as tolerated (8). One physician performed all clinical examinations, including modified Rodnan skin scores (mRSS).

Peripheral blood was obtained from venipuncture at three time points, referred to as timepoint 1 (*baseline* if treatment naïve or within two months of treatment initiation), timepoint 2 and timepoint 3 (**Supplementary Table 1**). Serum autoantibodies (ACA, RNAIII, and/or anti-topoisomerase I/Scl70) were measured by indirect immunofluorescence at Specialty Laboratories, Valencia, CA. Blood collection for research was performed at the time of clinically indicated testing. Patients were not involved in planning the study.

### Research participant data

Clinical variables recorded included patient age at blood collection, biological sex, SSc subtype (limited or diffuse cutaneous), skin disease trajectory [improved, stable, worsened as described (8)], blood collection date, treatment status (e.g., MMF or other immune suppression treatment), interstitial lung disease on high-resolution chest computed tomography, and pulmonary arterial hypertension (PAH) status (defined as a mean pulmonary artery pressure (mPAP) > 20mmHg and pulmonary vascular resistance (PVR) >2 WU on right heart catheterization) (**Table 1**).

**Table 1:** Systemic sclerosis patient study participants. Metrics include the patient ID in 4 numerical digits, age at study baseline, biological sex, SSc cutaneous disease subtype, anticentromere antibody (ACA), anti-RNA polymerase III (RNAIII) antibody, anti-topoisomerase I (Scl-70) antibody, presence of any SSc antibodies (ACA, RNAIII or Scl-70), antinuclear antibody (ANA), interstitial lung disease (ILD) defined as chest high-resolution computed tomography (HRCT) performed + or –, and evidence for ILD on HRCT + or -, pulmonary arterial hypertension (PAH) defined as right-heart catheterization performed + or – and mean pulmonary artery pressure (mPAP) >20mmHg with pulmonary capillary wedge pressure (PCWP) ≤15 mmHg, mycophenolate mofetil (MMF), and clinical skin improvement from baseline to study conclusion. (“+” indicates present, “NA” indicates results were not available because patient was not tested, blank indicates absence).

| Patient ID | Baseline Age | Biological Sex | SSc Subtype | ANA | SSc Antibodies | ANA | ACA | RNAIII | Scl70 | ILD | PAH | Skin Treatment | Skin Improvement |
| --- | --- | --- | --- | --- | --- | --- | --- | --- | --- | --- | --- | --- | --- |
| SScReg_1073 | 66 | F | Diffuse | + | + | + |  | + |  | +/+ | +/+ | No Treatment | Stable |
| SScReg_1469 | 49 | M | Diffuse | + | + | + | + |  |  | +/- | +/+ | No Treatment | Worse |
| SScReg_1736 | 42 | F | Diffuse | + | NA | + |  | NA |  | +/- | -/- | Other | Worse |
| SScReg_1753 | 33 | M | Diffuse | + | + | + |  | + |  | +/- | -/- | MMF | Improver |
| SScReg_1759 | 60 | F | Diffuse | + |  | + |  |  |  | +/+ | -/- | MMF | Stable |
| SScReg_1822 | 43 | F | Limited | + | + | + |  | + |  | +/- | -/- | MMF | Stable |
| SScReg_1823 | 61 | M | Limited | + | + | + | + |  |  | +/+ | -/- | No Treatment | Stable |
| SScReg_1832 | 41 | F | Limited | + |  | + |  |  |  | +/+ | -/- | MMF | Stable |
| SScReg_1837 | 30 | F | Diffuse | NA | + | NA | NA | + | NA | +/+ | -/- | MMF | Stable |
| SScReg_1858 | 48 | F | Diffuse | + | + | + |  |  | + | +/- | -/- | No Treatment | Worse |
| SScReg_1861 | 47 | F | Limited | + | + | + | + |  |  | +/- | -/- | MMF | Stable |
| SScReg_1886 | 58 | M | Diffuse | + | + | + |  | + |  | +/+ | -/- | MMF | Stable |
| SScReg_1891 | 51 | F | Diffuse | + | + | + |  | + |  | +/- | -/- | MMF | Worse |
| SScReg_1896 | 63 | F | Diffuse | + |  | + |  |  |  | +/+ | -/- | MMF | Improver |
| SScReg_1904 | 55 | F | Diffuse | + | + | + |  | + |  | +/- | -/- | MMF | Worse |
| SScReg_1906 | 70 | F | Diffuse | + | + | + |  | + |  | +/+ | -/- | Other | Stable |
| SScReg_1916 | 55 | F | Diffuse | + | + | + |  | + |  | +/- | +/+ | MMF | Worse |
| SScReg_1919 | 38 | F | Diffuse | + |  | + |  |  |  | -/NA | -/NA | MMF | Worse |

### CD4^+^ T-cell isolation and ATAC-seq library preparation

Lymphocyte isolation for ATAC-seq library preparation was performed as previously described (9). CD4^+^ T-cells were first enriched from whole blood using RosetteSep Human CD4^+^ T Cell Enrichment Cocktail (StemCell Technology) and cryopreserved in 10% DMSO/FBS for liquid nitrogen storage. Frozen cells were thawed and washed twice in 5% FBS in PBS 1X, filtered through 100 uM cell strainers, stained for viability with 7-AAD (BD Biosciences, 559925) and anti-human T-cell markers CD3 (Thermo Fisher Scientific, 11-0037-41) and CD4 (Tonbo Biosciences, 20-0048-T025). The BDFACS ARIA II with a 100 uM nozzle was used to FACS-isolate live 7-AAD^-^ CD3^+^ CD4^+^ T-lymphocytes for Omni-ATAC library prep (5). Sequencing library quality metrics are included in **Supplemental Table 2** and **Supplemental Fig. 1**.

### Mitochondrial mutation calling and patient samples consistency confirmation

Mitochondrial SNP profiling was used to confirm patient identity across timepoints using a previously described pipeline (10). Mitochondrial mutations were called using the GRCh37 reference from the 1000 Genomes Project and the mtDNA sequence rCRS. Reads aligned to the mitochondria reference genome were extracted from paired-end ATAC-seq fastq files aligned to the reference genome using BWA (11). Samtools (11) was used to convert mitochondrial sam files to bam files, sort bam files, and remove duplicated reads. Samtools mpileup generated the pileup file for each sample with option “-q 30 -Q 30”. A custom perl script was used to extract mutation information and filter low quality reads. High-confidence mitochondrial SNPs (allele frequency > 0.9, support reads > 3) were visualized by heatmap to confirm mitochondrial mutation consistency across samples from three timepoints for each patient **(Supplemental Fig. 1A**).

### ATAC-Seq data analysis

ATAC-seq data was processed and analyzed as previously described (5), with notable exceptions. Following adaptor trimming, reads were aligned to the hg38 genome using botwie2 (12) and filtered for duplicate reads and reads with low alignment score (<10). ATAC-seq peaks were annotated using GREAT (13) under the basal plus extension default setting.

Differential ATAC-seq peaks were called using negative binomial models from the R package DESeq2(14). Multiple hypothesis testing was controlled for using the Benjamini-Hochberg method to estimate false discovery rate (FDR (15). Peaks were defined as significant with absolute fold change larger than 1.5 and either FDR < 0.2 for sex and age factors or FDR < 0.3 for ACA and RNAIII comparisons.

Healthy control CD4+ T cell ATAC-seq data (GSE85853)(16) was downloaded from GEO and analyzed as described for SSc sample data. To avoid batch effect concerns, downloaded healthy control data were compared only with each other and not directly with the SSc samples from our study.

### CIBERSORT analysis

CD4+ T-cell subset proportions were inferred using CIBERSORT (17) and CD4 T cell ATAC-seq reference data (GSE118189 (18)). The overlap peak set between the SSc ATAC-seq peaks and reference peaks was generated using the Bedtools intersect module. The overlap peak set, mixture file, reference sample file, and phenotype files were generated according to the CIBERSORT manual. The R package edgeR (19) was used to normalize both Mixture count matrix and reference count matrix, and the subsequent matrices converted to log2CPM value. The signature file and the final CD4 T cell subtypes fraction matrix were then generated using CIBERSORT.

### GWAS enrichment and GO term analysis

GWAS enrichment analysis used SNPs indexed in autoimmune and control diseases from the European Bioinformatics Institute GWAS Catalog (https://www.ebi.ac.uk/gwas/docs/file-downloads) with Linkage Disequilibrium (LD) r^2^ > 0.8 (haploreg: http://archive.broadinstitute.org/mammals/haploreg/data/). The ucscTools liftOver module was used to lift SNP positions to the hg38 genome and numbers of SNPs within the SSc ATAC-seq peak regions were calculated by disease.

A random shuffle procedure was conducted 1000 times using the bedtools shuffle module to construct a null background from random peak sampling from which empirical p values were computed. The fold enrichment of GWAS analysis was calculated as the observed number of overlapping SNPs versus the mean random shuffled background.

SSc GWAS SNPs within our SSc ATAC-seq data were recorded as SSc ATAC-specific GWAS SNPs. Identities of associated SNPs were assigned by gene proximity and passed to Gprofiler (20) for functional gene set enrichment analysis.

### HiChIP data Analysis

HiChIP data for Naïve T cells, Th17 cells and Treg were downloaded from GEO (GSE101498(21)). Loops were called using FitHiChip (22) with 5kb bin, peak-to-all interaction type, loose background, and FDR < 0.01. LiftOver was used to convert the merged significant interaction files generated from the FitHiChIP pipeline from hg19 to hg38 and then to BigBed format. hg38-aligned files were then converted to BigBed format as described in the FitHiChIP online manual (https://ay-lab.github.io/FitHiChIP/usage/output.html) and visualized in UCSC web browser.

### Motif analysis

Homer (23) was used to conduct motif enrichment analysis using the intersecting regions between SSc ATAC-seq peaks and significant 5kb binned loops anchors called by FitHiChIP pipeline. Motifs with FDR < 0.05 were considered significant for each comparison.

### Statistical Analyses

All statistical analyses and quantification were conducted in R or GraphPad Prism as described. Statistical details of the experiments and assays, including exact “n” values, statistical tests, comparisons, and cutoffs are provided in the figure legends.

### Data Availability

All de novo ATAC-seq files and processed files generated for this study have been deposited in the Gene Expression Omnibus (GEO) under the accession identifier GSE163066.

## RESULTS

This study included 18 SSc patients (14 dcSSc) with 3 longitudinal timepoints. The median SSc disease duration between first Raynaud symptom and the initial visit was 16 months and median modified Rodnan skin score (mRSS) was 13.5. ACA, RNAIII, and Scl-70 antibodies were present in 17%, 50%, and 6% of patients, respectively. (**Supplementary Table 1, Table 1**)

### Longitudinal profiling of systemic sclerosis patients

Purified CD3^+^CD4^+^ T-lymphocytes were isolated from peripheral blood for ATAC library preparation (5) (**Figure 1A**). ATAC-seq library quality checks and mitochondrial SNP profiling to confirm patient longitudinal sample identity were performed prior to analyses (**Supplemental Table 2, Supplemental Figure 1**).

**Figure 1:**
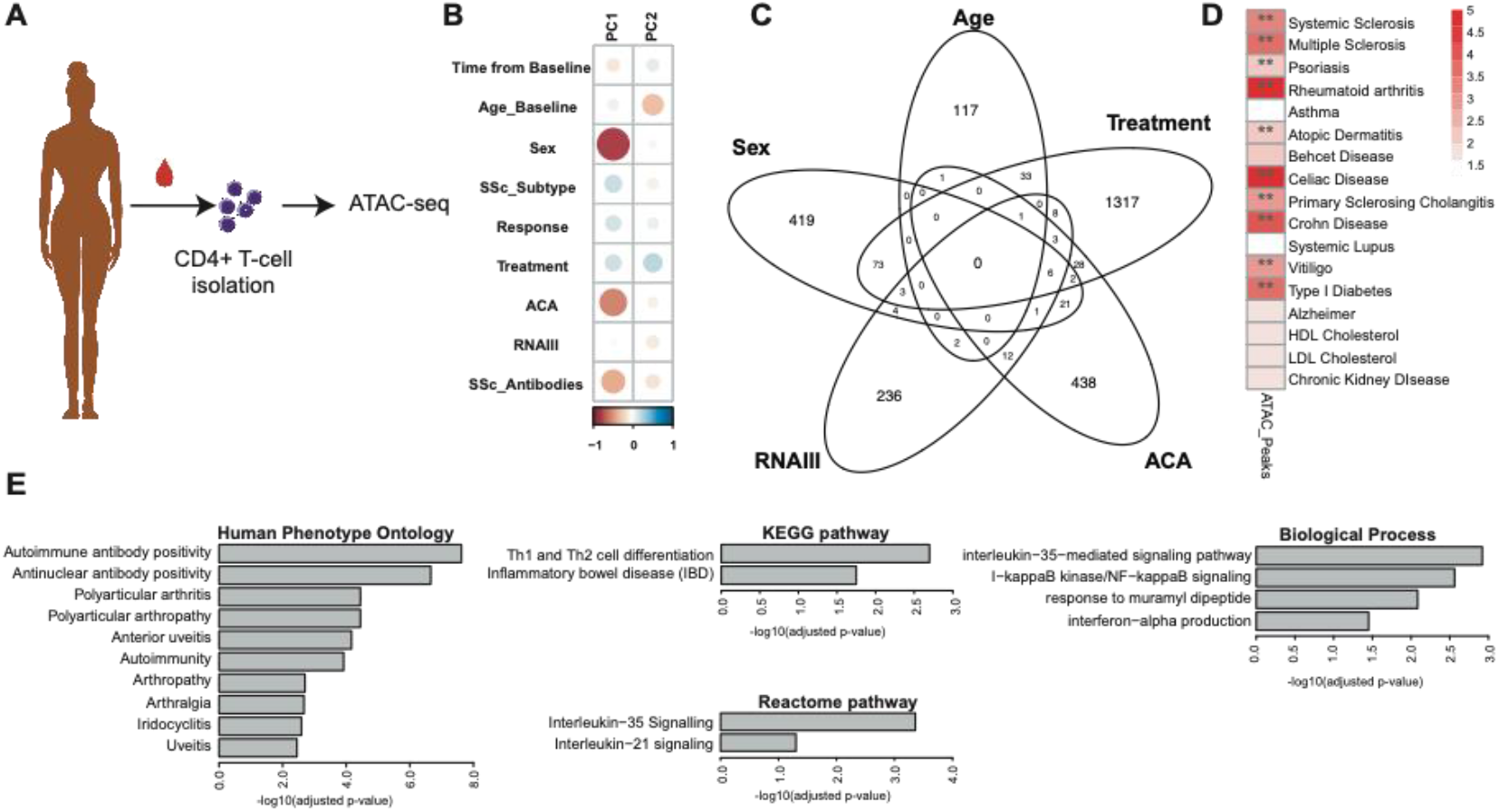
General characteristics of SSc study patients. **A** Diagram of sample collection from patients and ATAC-seq library preparation of CD4^+^ cells. **B** Spearman Correlation between PC1/PC2 and each contributing variable tested, color indicates spearman correlation value (from -1 red to 1 blue), size indicates strength of correlation (larger is stronger). **C** Differential peaks across all timepoints and samples for each of the main variables. **D** enrichment of GWAS SNPs in open chromatin regions from the SSc study samples to verified autoimmune disease-associated SNPs. Scale bar corresponds to fold enrichment of SNPs in diseases compared to the random sampling background (red=high enrichment, white = low enrichment), ** indicates FDR < 0.05 and fold enrichment > 2. **E** GO term and pathway analysis of SSc SNPs enriched ATAC peaks associated genes. ATAC-seq data analyzed using DESeq 2, statistics calculated using negative binomial modeling. N=18 patients across 3 timepoints each (see **Supplementary Tables 1** and **2**).

PCA performed on all sample ATAC-seq data identified sex, followed by SSc autoantibody status (specifically ACA positivity) and, to a lesser extent, age and treatment, contributed most to chromatin accessibility variance (**Figure 1B, Supplemental Figure 1E**). Minimal overlap was observed between peaks associated with different variables (**Figure 1C**), indicating independent contributions. GWAS-enrichment analysis identified significant overlap with multiple autoimmune disease loci, including SSc (**Figure 1D)**. Associated ontology from GWAS SNPs enriched in our dataset further highlighted autoimmune-related pathways and categories as well as SSc-specific disease-drivers, such as Th1/Th2 differentiation, IL-35 and IL-21 signaling, NFKB signaling, and IFN-alpha production (**Figure 1E**)(24).

### MMF treatment in SSc does not alter circulating CD4^+^ T-cell chromatin accessibility

To assess the potential impact of MMF treatment on T helper cells in SSc, we identified patients who donated a blood sample within two months of treatment initiation (baseline) as well as at two subsequent treatment timepoints (**Supplemental Table 1**). Comparison of the CD4^+^ T-cell ATAC-seq profile of eight MMF-treated, and four untreated patients over the three timepoints showed a stable, unchanging epigenetic pattern in CD4^+^ T-cells from MMF-treated patients and very few changes in untreated patients (**Figure 2A, 2C**). Similarly, CIBERSORT analysis showed MMF treatment had no significant impact on the relative proportions of T-cell subsets within the CD4^+^ population (**Figure 2D, Supplemental Figure 2**). Baseline differential peaks between the MMF-treated and untreated groups (**Figure 1C**) persisted over time, suggesting these differences reflect disease severity rather than MMF effect (**Figure 2B**). Based on these results, MMF treatment in SSc patients does not change the circulating CD4^+^ T-cell epigenome.

**Figure 2:**
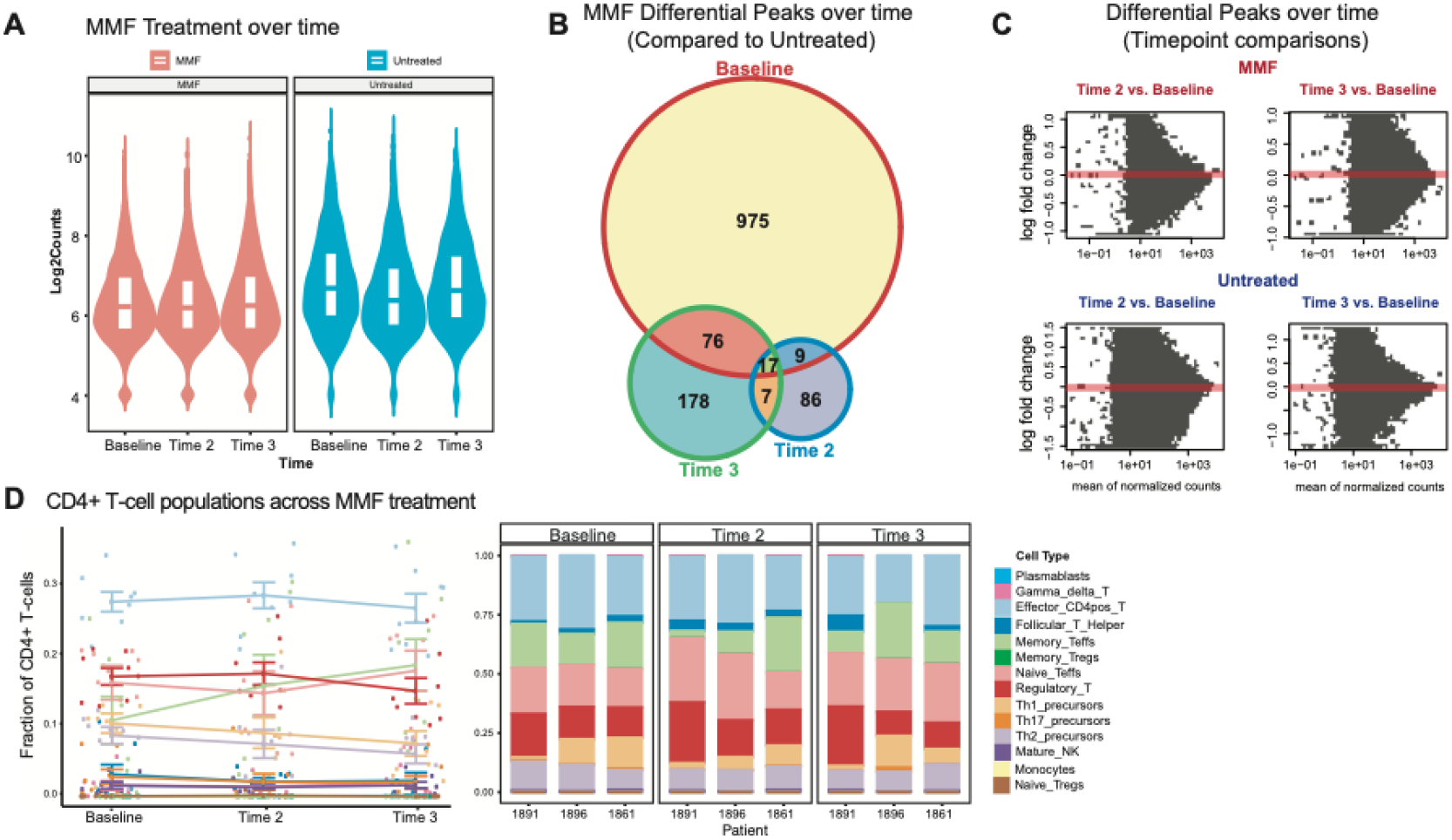
MMF treatment has no significant effect on CD4+ epigenome in SSc study patients. **A** Violin plot of differential ATAC peaks between MMF vs. healthy across all three timepoints in MMF-treated (red) and treatment naïve (blue) SSc patients. **B** Venn diagram of number of differential peaks in MMF-treated patients as compared to treatment naïve (untreated) across timepoints. **C** MA plots of differential peak comparisons between timepoints in both MMF-treatment and untreated groups. Red dots indicate significance. **D** CIBERSORT analysis of CD4^+^ T-cell subsets in MMF-treatment patients across time (composite of all patients by cell type, left, individual patient examples, right). N=10 MMF-treated patients, N=4 Treatment naïve patients, all across 3 timepoints each.

### Sex differences contribute the greatest variance in the circulating CD4^+^ T-cell regulome in SSc

The prevalence of most autoimmune diseases, particularly SSc, is higher in females than males(25). PCA results from our study show that the greatest variance arises from biological sex differences (**Figure 1B, Figure 3A**) with 82 of 203 differential peaks arising from the X chromosome in females with SSc, 4 and 80 of 323 mapping to the X and Y chromosomes, respectively, in males with SSc, and the remaining differential peaks to the autosomes (**Supplementary Figure 3A**). To focus the differential analyses exclusively on sex differences, we compared only SSc females and males at baseline (if treated, or at the first available timepoint if untreated) and identified clear segregation of gene regions more accessible in females with SSc distinct from those in males with SSc (**Figure 3B, 3D**).

**Figure 3:**
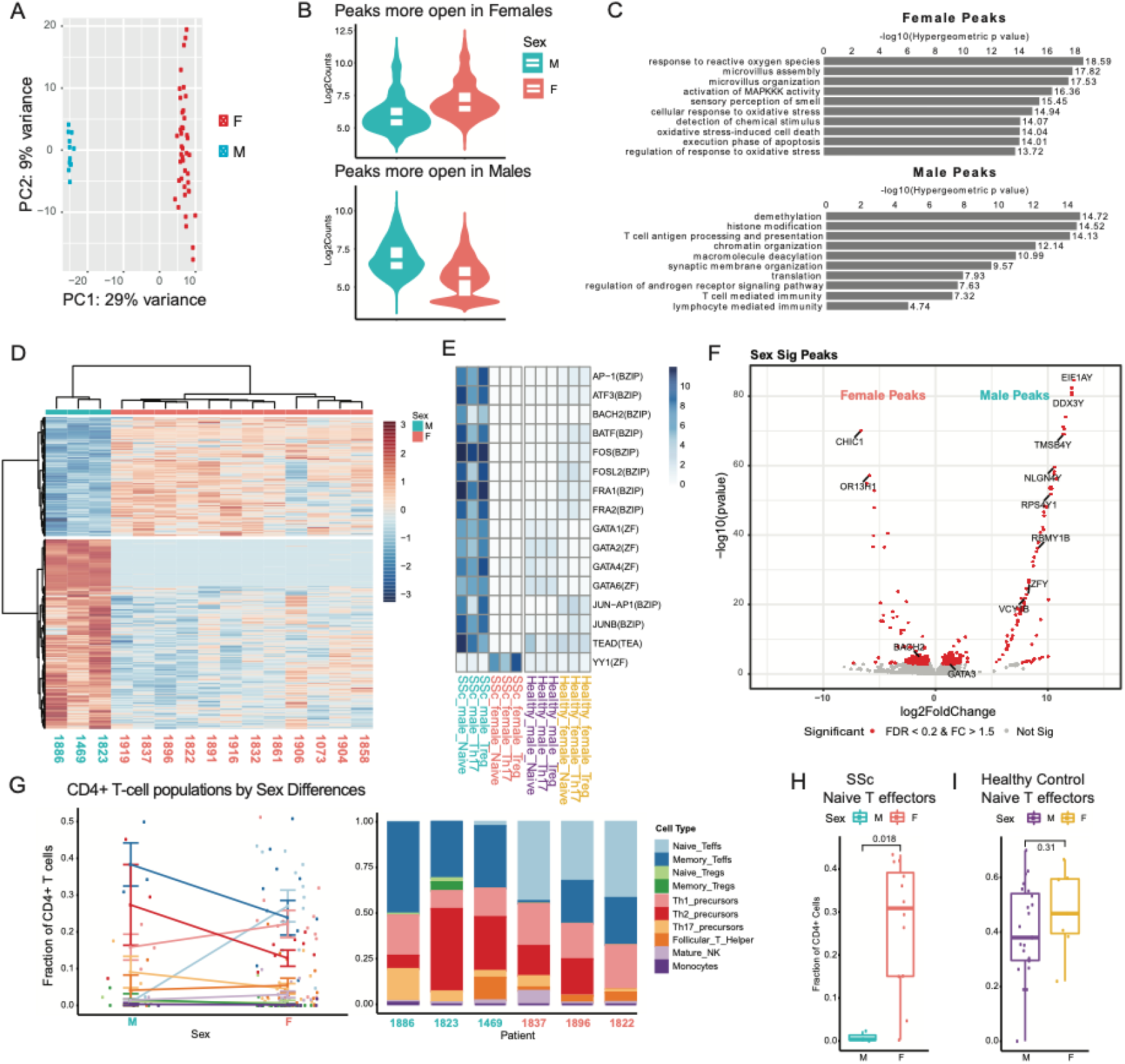
Greatest variance in SSc study patients arises from sex differences. **A** PCA analysis of all patients and all timepoints grouped by sex (N=18, across 3 timepoints each). **B** Violin plots depicting peaks more accessible in females (red) and in males (blue) (N=18 patients, across 3 timepoints each). **C** GREAT predicted functions of cis-regulatory regions associated with peaks accessible in SSc females and males. P-values calculated using Fisher’s Exact Test (N=18 patients, across 3 timepoints each). **D** Heatmap of differential ATAC-seq peaks between male and female SSc study patients at treatment-naïve, baseline timepoint (SSc-1736, -1753, and -1759 excluded because they had >2 months of treatment at the 1^st^ timepoint) grouped by sex (red = female, blue = male), scale bar indicates z-score of accessibility (red is more accessible and blue is less accessible, N=3 males and N=12 female SSc patients shown). P-values calculated using negative binomial models in DESeq2. **E** Heatmap of motif enrichment of the genomic region from overlay of significant ATAC-seq peaks with HiChIP anchors, scale bar indicates significance of enrichment (values correspond to -log10(p-value), darker blue is more significant) (SSc N=15 patients; Healthy controls N=10 donors, 31 total samples, downloaded from GEO GSE85853). **F** Volcano plot of differential peaks with FDR < 0.2 and absolute fold change larger than 1.5 highlighted in red, significance calculated using negative binomial models in DEseq2. **G** CIBERSORT analysis of CD4^+^ T-cell subsets in male and female SSc patients (left panel: composite of N=15 patients by cell type; right panel; individual patient examples). **H** Percentage of Naïve T effectors grouped by sex in SSc study patients (SSc N=15 patients at baseline timepoint) calculated by CIBERSORT **I** Percentage of Naïve T effectors in general population control patients calculated by CIBERSORT (31 total samples from N=7 males and N=3 females, GEO GSE85853).

To provide a comparison basis for sex-related differences unrelated to disease, healthy control data from published CD4^+^ T-cell ATAC data (9) were analyzed separately from our SSc data. Interestingly, there were fewer differential peaks arising from sex differences in the general population (Females = 54, Males = 154) and a larger fraction of differential peaks arose from the sex chromosomes (Female X = 77.8%, Male Y = 39.0%) than in SSc males and females (**Supplemental Figure 3B**). This disparity suggests that factors other than the sex chromosomes may drive the sex differences observed in SSc.

GREAT analysis of the differential peaks identified high prevalence of multiple immune-related pathways. Strikingly, T-lymphocyte-related programs associated to peaks more accessible in SSc males than in SSc females (**Figure 3C**). Specific to SSc males, differentially accessible ATAC-seq peak regions mapped to significant motif enrichment of genes involved in immune response and increased T-cell activation (AP-1, FOS, JUN, BATF) (**Figure 3E**) and the Th2-activator GATA3 (**Figure 3F)**. CIBERSORT analysis of the CD4^+^ T-cell subtypes also revealed a trend of a higher proportion of pathogenic Th2 cells in SSc males (**Supplemental Figure 3E**). Conversely, SSc females exhibited a significantly higher proportion of the more undifferentiated naïve T effector cells (**Figure 3G, 3H**). These sex differences were not present in the general population (**Figure 3I, Supplemental Figure 3F**), suggesting SSc-specific disease divergences in differentiation and homeostasis of T effector cells between sexes.

While many of the same predicted cis-regulatory functions appeared in both healthy and SSc females, neither group showed motif enrichment of the SSc male-associated chromatin regions (**Figure 3C, Supplementary Figure 3C, Figure 3E**). Since biological females are more prone to chronic autoimmunity than males, this result is unsurprising. Additionally, males who develop SSc tend to have much more severe disease than females (25, 26), further supporting the hypothesis that more changes in immune-related expression are required to push males from homeostasis to disease.

### Aging has limited impact on chromatin accessibility in SSc peripheral T-cells

Although all patients in our SSc study are adults, age is a potential differential contributing factor that arises when sex-related differences are excluded from the comparison (**Figure 1B, Supplemental Figure 1E**). In contrast to sex, age-related differential peaks were more evenly distributed across both autosomes and sex chromosomes (**Supplemental Figure 3G**). To avoid potentially confounding factors (e.g., drug metabolism differences), only treatment-naïve or samples at baseline were included for age-specific analyses (**Supplemental Figure 4**).

We observed a trend of increased chromatin accessibility in circulating CD4^+^ T-cells with advancing age (**Supplemental Figure 4A**). Heat maps of accessible peaks revealed two distinct clusters distinguishing younger versus older patients (**Supplemental Figure 4B**). Accessible genes in older patients were related to immune activity pathways, such as the JUN-AP1 and BACH2-BATF motifs, interferon response factor IRF2 and T-cell proliferation regulator TNFSF8 (**Supplemental Figure 4C, D**). CIBSERSORT assignment of the CD4^+^ T-cell subsets based on ATAC-seq signatures did not show any statistically significant or consistent trends between the three age categories (**Supplemental Figure 4F**). These findings align with previous findings of immune system aging (27) and suggest that the effects of aging on CD4^+^ T-cell chromatin accessibility is not a major mechanism of SSc pathobiology.

**Figure 4:**
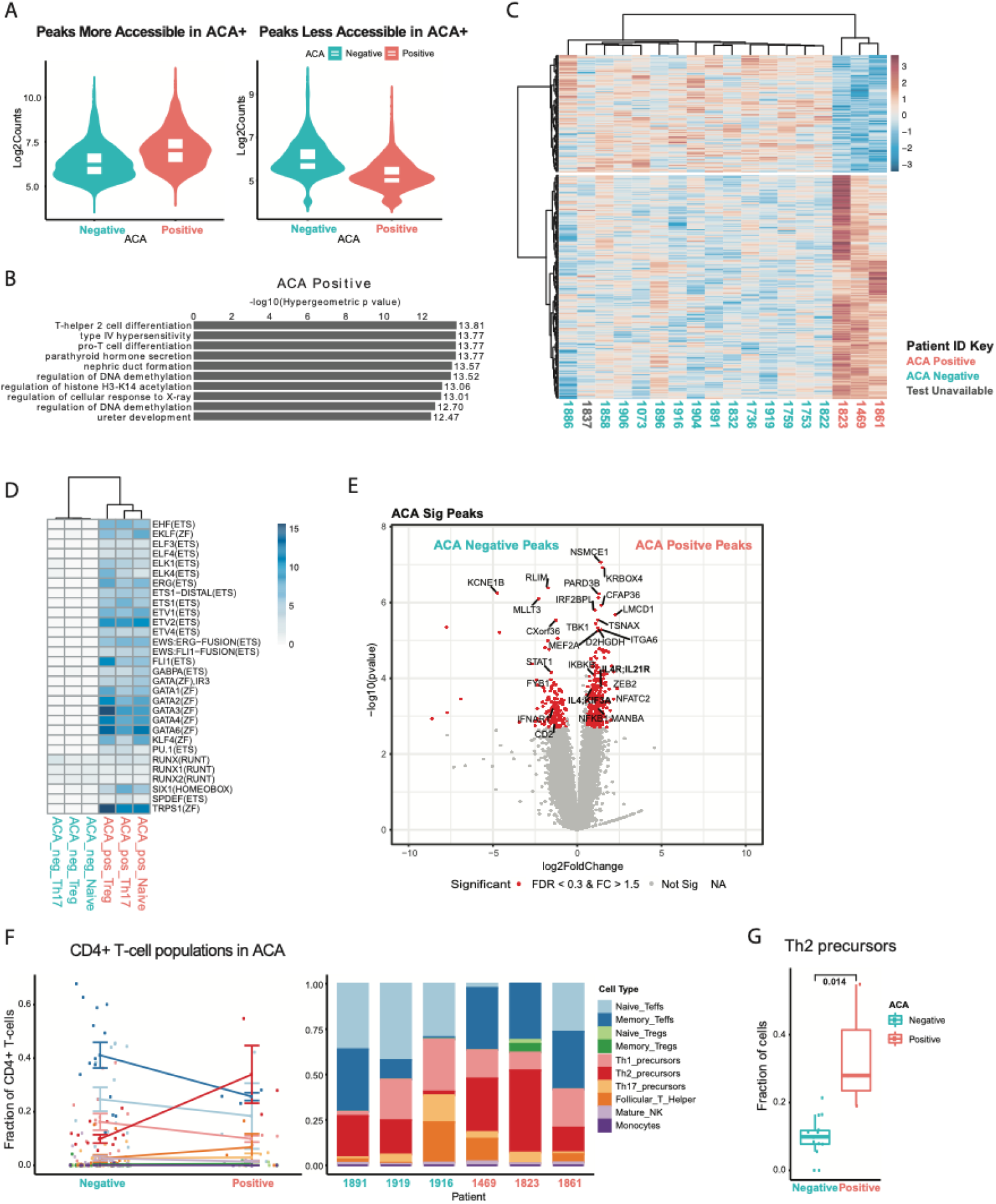
ACA-positive patients present with more Th2 cells. **A** Violin plots depicting peaks more accessible in ACA-positive (red) and in ACA-negative (blue) patients. **B** GREAT predicted functions of cis-regulatory regions associated with peaks accessible in ACA-positive patients (GREAT analysis for ACA-negative patients not significant). Statistics calculated using Fisher’s Exact Test, N=18 patients, 3 timepoints each. **C** Heatmap of differential ATAC-seq peaks of SSc study patients at latest timepoint (N=18 patients, timepoint 3 shown) available grouped by ACA-positive (red) and negative (blue). Grey indicates status not reported. Scale bar indicates z-score of accessibility (red is more accessible, and blue is less accessible) **D** Heatmap of motif enrichment of the genomic region from overlay of differential ATAC-seq peaks with HiChIP anchors, scale bar indicates significance of enrichment (values correspond to -log10(p-value), darker blue is more enriched and significant. N=18 patients, 3 timepoints each. **E** Volcano plot of differential peaks with FDR < 0.3 and absolute fold change larger than 1.5 highlighted in red (N=18 patients, 3 timepoints each). **F** CIBERSORT analysis of CD4^+^ T-cell subsets in confirmed ACA-positive and negative patients (SSc-1837 excluded, composite of all patients by cell type, left, individual patient examples, right). **G** Percentage of Th2 precursor cells grouped by ACA-status in SSc study patients (N=17: 14 negative and 3 positive at timepoint 3) calculated by CIBERSORT (Wilcoxon Rank Sum Test).

### Autoantibody profiles correlate with Th2-associated pathways

Serum autoantibodies were the most significant non-sex related factor affecting CD4^+^ T-cell chromatin accessibility in SSc (**Figure 1B, Supplemental Figure 1E**). Because Principal Components Analysis (PCA) showed that ACA positivity individually had larger variation than all three SSc autoantibodies grouped together (i.e., expressed autoantibodies vs. none) (**Figure 1B**) we focused on the individual autoantibody groups. We used the 3rd timepoint because the most pronounced peak differences were observed at the latest (3^rd^) time point collected, potentially because of prolonged disease progression. All patients with available ACA and RNAIII antibody data were included as there was little overlap of total differential peaks across timepoints between each of the comparison factors (**Figure 1C, Table 1**).

ACA and RNAIII antibodies are mutually exclusive in > 95% of patients (28) and associate with different clinical outcomes: PAH and scleroderma renal crisis most commonly present in ACA and RNAIII-positive SSc patients, respectively (29). Given these clinical attributes, the lack of a significant number of Scl70 patients, and the presence of ANA in nearly every patient, we focused on differences arising from ACA and RNAIII expression (**Table 1**). A point of interest comes from patient 1837, who was not included in our ACA analysis due to lack of an ACA confirmatory test. However, since patient 1837 has confirmed expression of RNAIII autoantibodies, we can reasonably infer the absence of ACA(28). Furthermore, the ATAC-seq profile of subject 1837 clustered independently with the ACA negative patients, suggesting that chromatin accessibility profiles may predict autoantibody expression.

There was a clear distribution pattern of accessible DNA elements in CD4^+^ T-cells in positive-compared to negative-ACA patients (**Figure 4A, C**). GREAT analysis of cis-regulatory functions of differential peaks revealed that more accessible peaks corresponded to Th2, cell-mediated immune responses, and T-cell differentiation in ACA+ patients (**Figure 4B**). Overlap of differential peaks with Hi-ChIP anchor sites in CD4^+^ T-cell subsets showed enrichment of multiple motifs corresponding to immune cell function and autoimmunity (ETS family) (30), T-cell development (FLI1), Th2 differentiation (GATA3) as well as other immune cell programs (PU.1) that are significantly present in ACA+ T-cells but not ACA-T-cells (**Figure 4D**). Regions of genomic accessibility corresponding to the pro-inflammatory transcription regulator, NFKB1, and Th2 genes, IL4, IL4R, and IL21R, associated with ACA+ (**Figure 4E**). Additionally, CIBERSORT deconvolution of CD4+ T-cell subsets detected significant compositional differences characterized by higher Th2 proportions in ACA+ compared to ACA-patients (**Figure 4F, 4G**).

Next, we sought to determine the relationship between peaks in patients with different serum autoantibodies. As in the ACA comparison group, a clear distribution pattern distinguished RNAIII+ and RNAIII-genomic accessibility in CD4^+^ T-cells (**Figure 5A, C**). Upon closer examination of genomic regions and associated pathways, there were clear similarities shared between ACA- and RNAIII+ patients even though not all ACA-patients are also RNAIII+, as some patients were negative for both autoantibodies (ACA-/RNAIII-). As in the ACA+ group, a subset of RNAIII-patient cells showed greater and more differential peaks compared to RNAIII+ associated peaks (**Figure 5A**). These two different peak sets (RNAIII+ and RNAIII-) gave rise to two distinct clusters on the Z-score heatmap (**Figure 5C**).

**Figure 5:**
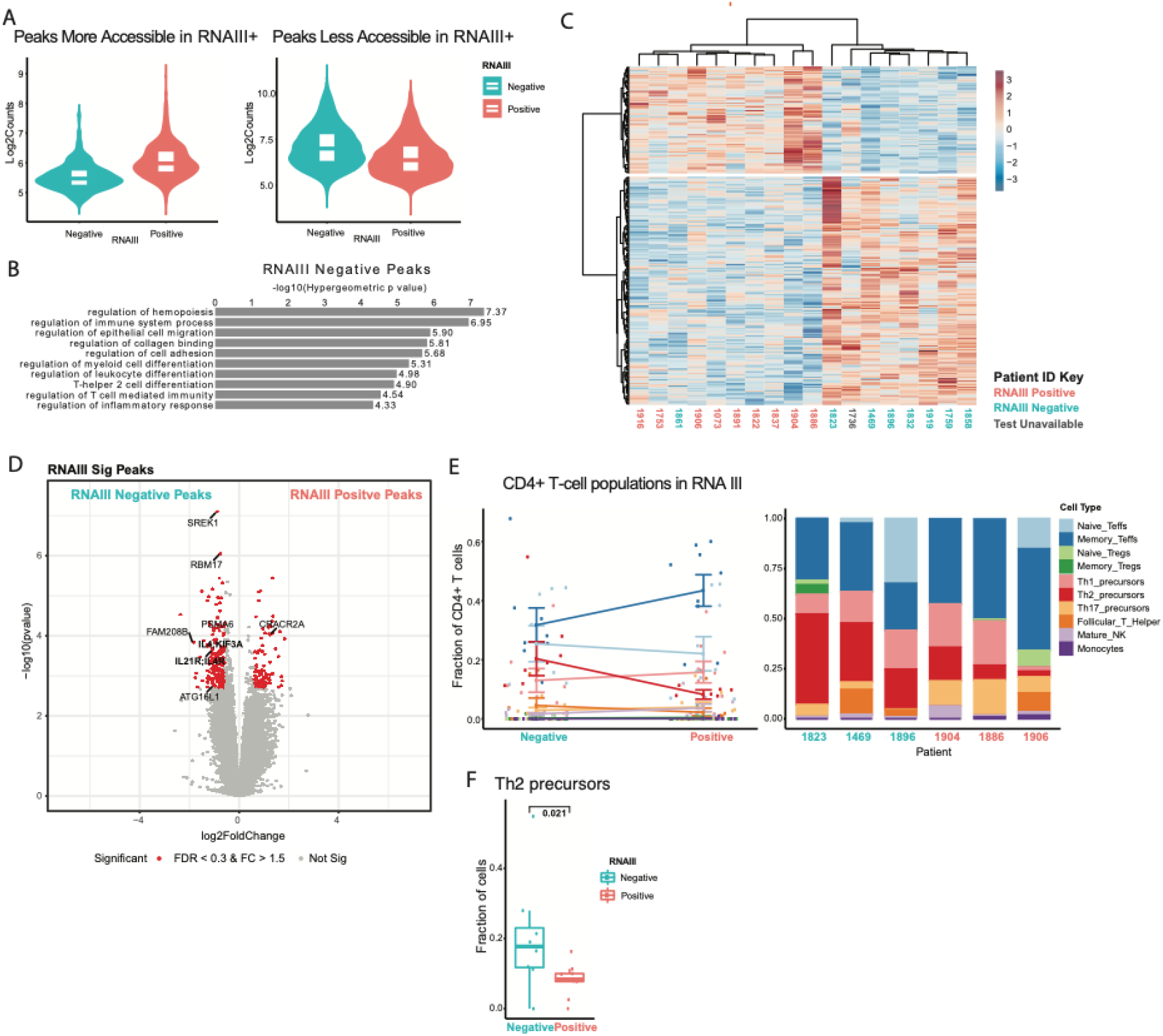
RNAIII-positive patients present with fewer Th2 cells. **A** Violin plots depicting peaks more accessible in RNAIII-positive (red) and in RNAIII-negative (blue) patients. **B** GREAT predicted functions of cis-regulatory regions associated with peaks accessible in RNAIII-negative patients (GREAT analysis for RNAIII-positive patients not significant). **C** Heatmap of differential ATAC-seq peaks of SSc study patients at latest timepoint (timepoint 3, N=18) available grouped by RNAIII-positive (red) and negative (blue). Grey indicates status not reported. Scale bar indicates z-score of accessibility (red is more accessible, and blue is less accessible) **D** Volcano plot of differential peaks with FDR < 0.3 and absolute fold change larger than 1.5 highlighted in red. **E** CIBERSORT analysis of CD4^+^ T-cell subsets in confirmed RNAIII-positive and negative patients (SSc-1736 excluded, composite of all patients by SSc study patients calculated by CIBERSORT (Wilcoxon Rank Sum test, N=17: 9 positive and 8 negative at cell type, left, individual patient examples, right). **F** Percentage of Th2 precursor cells grouped by RNAIII-status in timepoint 3).

GREAT analysis further revealed that CD4+ T-cells from ACA+ and RNAIII-patients overlapped across many immunity and T-cell related pathways, particularly Th2 differentiation (**Figure 5B**), and significant RNAIII-differential peak regions highlighted Th2-related genes such as IL4, IL4R, and IL21R (**Figure 5D**). These attributes were corroborated with CIBERSORT breakdown of the CD4^+^ T-cell subset that highlighted the significant difference in Th2 composition in RNAIII+ and RNAIII-cohorts. Both ACA+ and RNAIII-groups contained higher Th2 percentages when compared to ACA-/RNAIII+ groups (**Figure E, F**). Additionally, most differential peaks came from the ACA+ and RNAIII-groups while ACA- and RNAIII+ contributed fewer differential peaks (**Supplemental Figure 5A, B**), suggesting that major autoantibody-related differences primarily arise from ACA+ and/or RNAIII-carriage status.

### Anticentromere antibodies may be predictive of Th2-mediated fibrosis in SSc

We observed that CD4^+^ T-lymphocytes in SSc patients that are ACA+ and/or RNAIII-evinced significantly higher proportions of Th2 cells and related motifs and regulatory pathways. There were 44 total genes in the overlap of differential peaks associated with both ACA+ and RNAIII-, of which 19 were shared with SSc males, and featured multiple Th2-related genes, such as *IL4* and *IL21R* (ACA+ and RNAIII-only) and *GATA3* and *IL4R* (shared with SSc males) (**Figure 6A**). Here, we focus on the *IL-4-KIF3A* and *IL4R-IL21R* regions, which associated with ACA+ and RNAIII-in our analyses (**Figure 4E, Figure 5D**).

**Figure 6:**
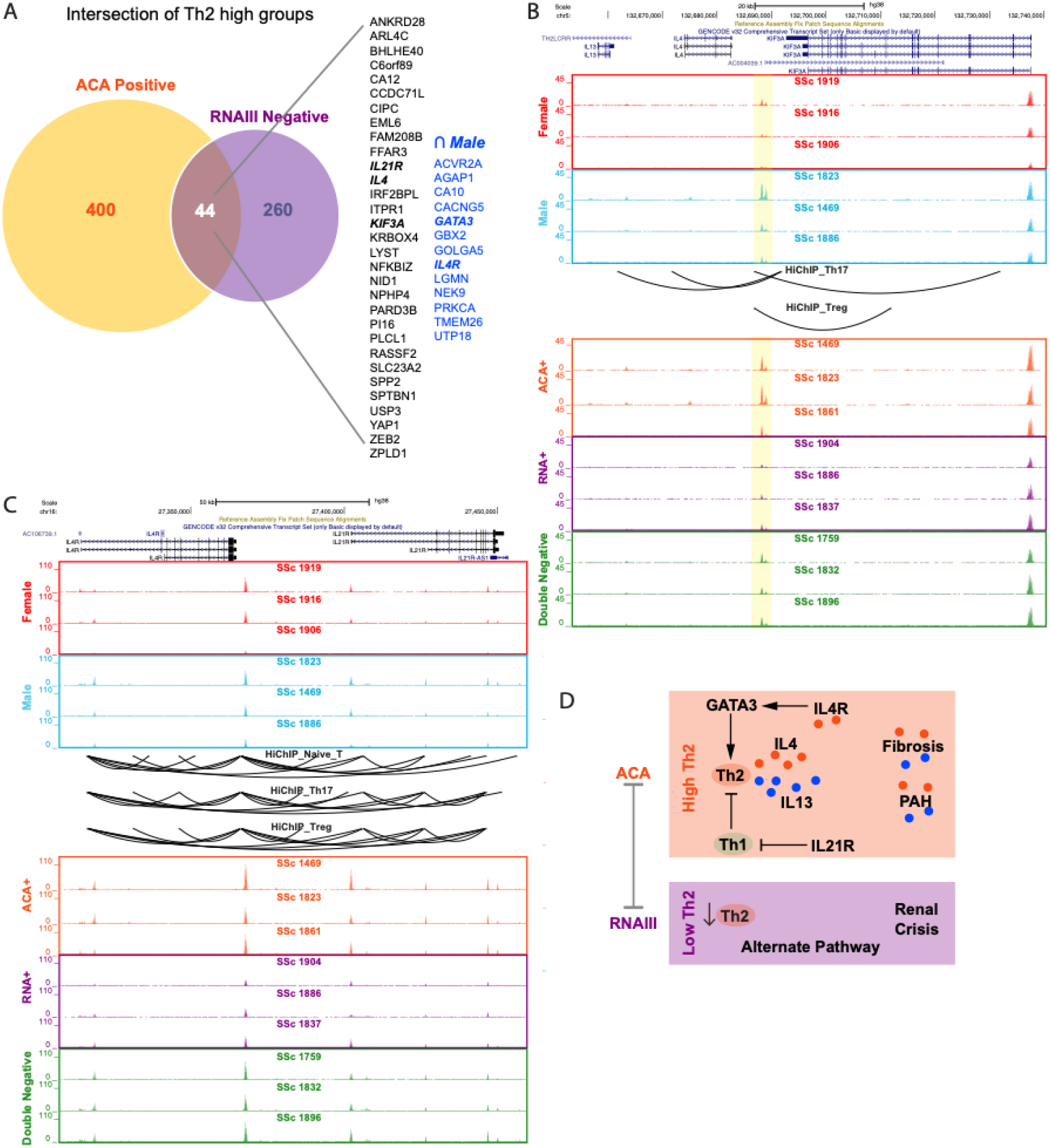
Interplay between autoantibody groups and Th2 pathways. **A** Venn diagram of accessible differential peaks in ACA-positive and RNA-negative SSc study patients (N=18 patients, timepoint 3). The genes within the intersection of the two autoantibody groups are listed and the intersection with male peaks are in blue. Bold italic indicates Th2-related genes. Representative ATAC-seq tracks for **B** the IL4;KIF3A and **C** the IL4R;IL21R genomic regions. Representative tracks for females at baseline (red), males at baseline (blue), ACA-positive at timepoint 3 (orange), RNA-III positive at timepoint 3 (purple), and ACA/RNAIII dual-negative at timepoint 3 (green) with HiChIP interaction loops from Naïve T-cell, Th17 and Tregs (black loops). The predicted enhancer for IL-13, IL-4, and KIF3A is highlighted in yellow. **D** Proposed model of relation between autoantibody expression and Th2 activation.

Th2 cells produce both IL-4 and IL-13 to stimulate fibrosis. The *IL-4* gene locus is in close genomic proximity to both *IL-13* and *KIF3A* (**Figure 6B**), another gene shared by the Th2-high groups (ACA+, RNAIII-). KIF3A is a ciliary protein important in myofibroblast development in SSc (31, 32). The expected trends for accessibility in the *IL-4-KIF3A* genomic interval was reflected across all three comparison groups (females vs. males, ACA+ vs. ACA- and RNAIII+ vs. RNAIII-) in relation to predicted Th2 cell proportions: peaks were larger in males compared to females, and significantly larger in ACA+ compared to RNAIII+. We also identified an active enhancer located between *IL-13* and *KIF3A* that is predicted through HiChIP to interact with *IL-4, IL-13*, and *KIF3A* in Th17 and Treg cells (**Figure 6B**). The peak levels of this putative enhancer reflect predicted Th2 activity levels.

Examination of *IL4R* locus in the *IL4R-IL21R* genomic region showed a similar accessibility pattern as the *IL4* region with higher accessibility peaks in samples with higher proportions of Th2 (**Figure 6C**). *IL21R* neighbors *IL4R* and also displays more chromatin accessibility in the Th2-high ACA+ CD4^+^ T-cells compared to the Th2-low RNAIII+ CD4^+^ T-cells (**Figure 6C**). IL21R is the receptor for IL-21, a potent cytokine secreted by both pathogenic Th2 and Th17 cells that promotes Th17 differentiation, inhibits Th1 differentiation, and drives fibrosis in diseases of chronic skin inflammation as well as pulmonary fibrosis in SSc(33-38). Interestingly, patients without either ACA or RNAIII antibodies (double negative) displayed accessibility levels intermediate between the higher ACA+ and lower RNAIII+ peak levels in both the *IL13-KIF3A* and *IL4R-IL21R* regions (**Figure 6B, C**), suggesting that expression of either autoantibody modulates Th2-related genomic accessibility, and potentially expression, patterns.

Our results point to an association between ACA+ and/or RNAIII-autoantibody status and higher Th2 prevalence and Th2-cytokine activity in SSc patients. Taken together with previous studies that identify Th2 cells as a driving force behind fibrosis in SSc (39, 40), the higher Th2 cell proportions and activity signatures detected in our study’s ACA+ and RNAIII-patients suggest autoantibody profiles impact circulating cellular pathway activation in SSc patients and may promote the differences observed in clinical phenotypes.

## DISCUSSION

Because autoimmunity and immune cell perturbations underlie SSc pathogenesis, we conducted a longitudinal study to comprehensively profile the chromatin landscapes of circulating CD4+ cells in SSc patients. Differences in CD4+ T-cell chromatin accessibility were associated with biological sex but not with age or MMF treatment. Importantly, we identify a novel link between anti-ACA+ serum autoantibodies and Th2-skewed immune responses, offering critical mechanistic insight into the clinically observed association between ACA and fibrotic complications such as PAH (28).

In prior studies, Th2 cytokines IL-4 and IL-13 are both detected at higher levels in SSc patient sera (41, 42), and have been implicated as drivers of PAH (43-46). Results from the tight skin mouse model, IL-4 has been strongly linked to dermal fibrosis: IL-4 ^-/-^ mice display less skin fibrosis (47) and addition of anti-IL-4 antibody reduces collagen production in IL-4 stimulated tight skin dermal cells (48). CD8+ production of IL-13 is directly associated with dermal fibrosis in SSc patients (49, 50). Thus, our data correlating ACA+ but not RNAIII+ to increased chromatin accessibility to the *IL13-IL4* genomic loci, underscore the relevance of autoantibody profiling especially when integrated with tracking of Th2-cytokines and cellular fractions.

MMF is used in autoimmune diseases for its immunomodulatory effects. However, we showed that MMF did not alter the chromatin state of circulating CD4+ T-cells over the study time course (**Figure 2**). It is possible that MMF works in other tissues or immune compartments (*e*.*g*., macrophages (3)) or requires longer duration to show effects. Our results suggest that targeted therapies to modulate Th2 activity can potentially be a specific and effective treatment option for ACA+ and/or for male patients with high levels of Th2-cytokines. Ongoing trials of IL-4/IL-13 blockade (42) further support this approach: (1) dupilumab (DB12159), an anti-IL-4 and IL-13 monoclonal antibody used to treat atopic dermatitis in Phase II trials for localized SSc (trial identifier: NCT04200755, 2019-002036-90, Uni-Koeln-3815) and (2) romilkimab (SAR156597), an IL-4 and IL-13 neutralizing IgG bi-specific antibody that appears to significantly improve skin fibrosis based upon the results of an early Phase II trial in early dcSSc patients (41). Given the recent trend towards adding immunomodulatory therapies, such as rituximab, to PAH-specific therapies in SSc-PAH (51) (Clinicaltrials.gov identifier: NCT01086540) it will be useful to explore if targeting IL-4/IL-13 in SSc-PAH patients, particularly those who are ACA+, is a more direct approach than adjunctive therapy. Moving forward, future trials of targeted therapies should include SSc autoantibody expression for consideration.

Herein, we have demonstrated that bulk ATAC-seq is a powerful and sensitive tool capable of high-fidelity detection of clinically meaningful epigenetic heterogeneity in SSc linked to sex and autoantibody status. From our pathway analysis, we propose that future clinical studies involving ACA+ SSc patients and anti-IL-4 and anti-IL13 therapy should be conducted. Future experiments will include SSc patients with other autoantibodies to identify pathways related to RNAIII+ and Scl70+, and involve other immune cell types including B- and myeloid cells. Additionally, as single cell ATAC-seq becomes increasingly cost-effective and commercially standardized, future studies should include chromatin profiles as an additional impactful SSc clinical phenotype.

## ACKNOWLEDGEMENTS

We thank members of the Chang lab and Dr. Oliver Distler for discussion.

## KEY MESSAGES

- **What is already known on this topic:** A biological female sex bias for acquiring SSc and a male sex bias for more severe SSc as well as an association with specific serum autoantibodies are known SSc features, but the precise immune cell subsets and regulatory programs involved in pathogenesis are not well understood and longitudinal studies to investigate the role of chromatin landscapes using ATAC-Seq in circulating CD4+ T cells is unexplored.
- **What this study adds:** ATAC-seq analysis of CD4^+^ lymphocytes was sufficient to consistently group participants by sex and autoantibody profile. Serum anticentromere (ACA) antibodies were associated with elevated T helper 2 (Th2) cell proportions and increased chromatin accessibility at gene regions encoding fibrosis-driving Th2 cytokines IL-4, IL-13, and the IL-4 receptor.
- **How this study might affect research, practice or policy:** SSc patients with anticentromere antibodies may be most likely to improve in response to drugs targeting Th2 cytokines. Inclusion of ATAC-seq profiling in clinical screens and registries may add significant value at low cost to help stratify disease and inform predictive models.

## AUTHOR CONTRIBUTIONS

Conceptualization: H.Y. Chang, D.R. Dou, and M.E. Hinchcliff; Methodology and Investigation: K. Aren, M. Carns, D.R. Dou, M. Hinchcliff, R. Li, L.C. Zaba, and Y. Zhao, N Field, A Peltekian; Data Curation: M Carns, K Aren, N Field, A Peltekian; Project Curation: K Aren and M Carns; Data Analysis: D.R. Dou, Y. Zhao, and B. Abe; Writing: B. Abe, H.Y. Chang, L.S. Chung, D.R. Dou, M. Hinchcliff, and Y. Zhao; Funding Acquisition: H.Y. Chang and M. Hinchcliff; Resources: H.Y. Chang and M. Hinchcliff; Supervision: H.Y. Chang and M. Hinchcliff; Writing and editing manuscript: H.Y. Chang, L.S. Chung, D.R. Dou, M. Hinchcliff, and Y. Zhao.

## DATA STATEMENT

All new ATAC-seq files and processed files generated for this study have been deposited in the Gene Expression Omnibus (GEO) under the accession identifier GSE163066. This new dataset is private and will be released to the public upon publication. Reviewers will be given a token for access.

Prior published HiChip (GSE101498 (77)) and healthy control ATAC-seq datasets can be accessed from GEO.

## DATA LINKING

https://www.ncbi.nlm.nih.gov/geo/query/acc.cgi?acc=GSE163066

## MATERIALS & CORRESPONDENCE

Further information and requests for resources and reagents should be directed to and will be fulfilled by Howard Y. Chang or the co-lead contact, Diana R. Dou.Queries about the clinical cohort in this study should be directed to the co-lead contact, Monique Hinchcliff.

## SUPPLEMENTAL TABLES

**Supplemental Table 1**: *Patient Timepoint Table*. Timepoint information (calculated in 0.5-month intervals) and baseline MRSS information for each patient.

**Supplemental Table 2:** *QC Table*. Quality Check metrics of ATAC-seq libraries for SSc study.

## SUPPLEMENTAL FIGURES

**Supplemental Figure 1:**
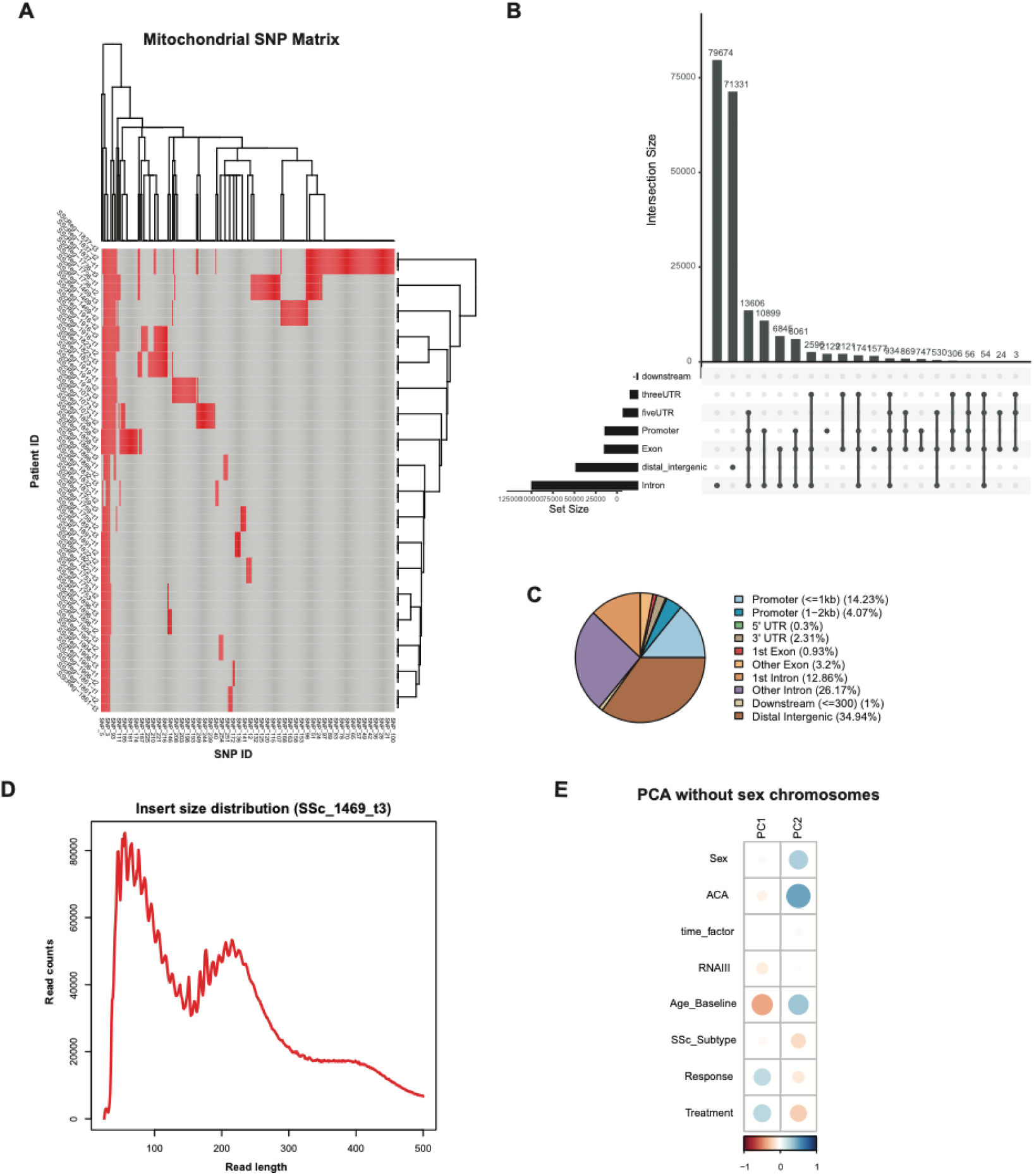
Quality check of ATAC-seq libraries for SSc study. **A** Matrix of mitochondrial SNPs for every patient at every timepoint to confirm veracity of patient ID across the study. **B** Graph of ATAC-seq peak distribution by number of peaks. **C** Venn diagram of ATAC-seq peak distribution by percentages. **D** Representative plot of ATAC-seq library fragment length distribution (SSc_1469_t3). **E** PCA plots of contributing variables in the study, excluding peaks from the sex chromosomes (autosomes only). N=18 patients, 3 timepoints each

**Supplemental Figure 2:**
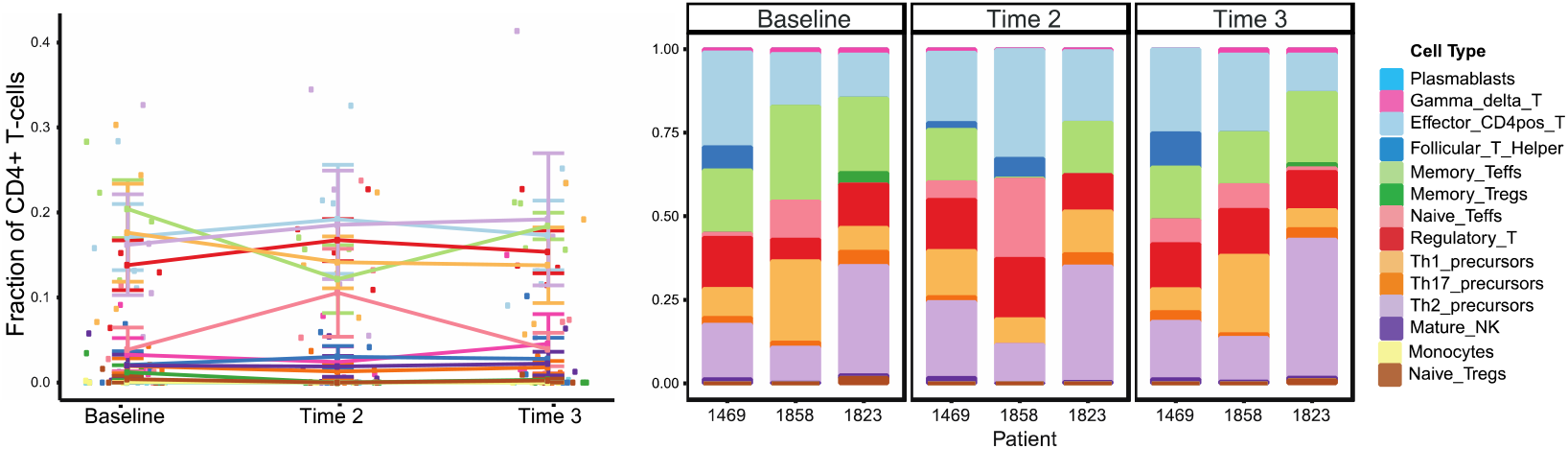
CIBERSORT analysis of CD4^+^ T-cell subsets in treatment naïve SSc patients across 3 timepoints. Left panel: composite of all 4 treatment naïve patients by cell type; Right panel: representative plots for 3 individual treatment naïve patient examples. Statistics calculated using the Wilcoxon Rank Sum Test, mean + SEM bars shown.

**Supplemental Figure 3:**
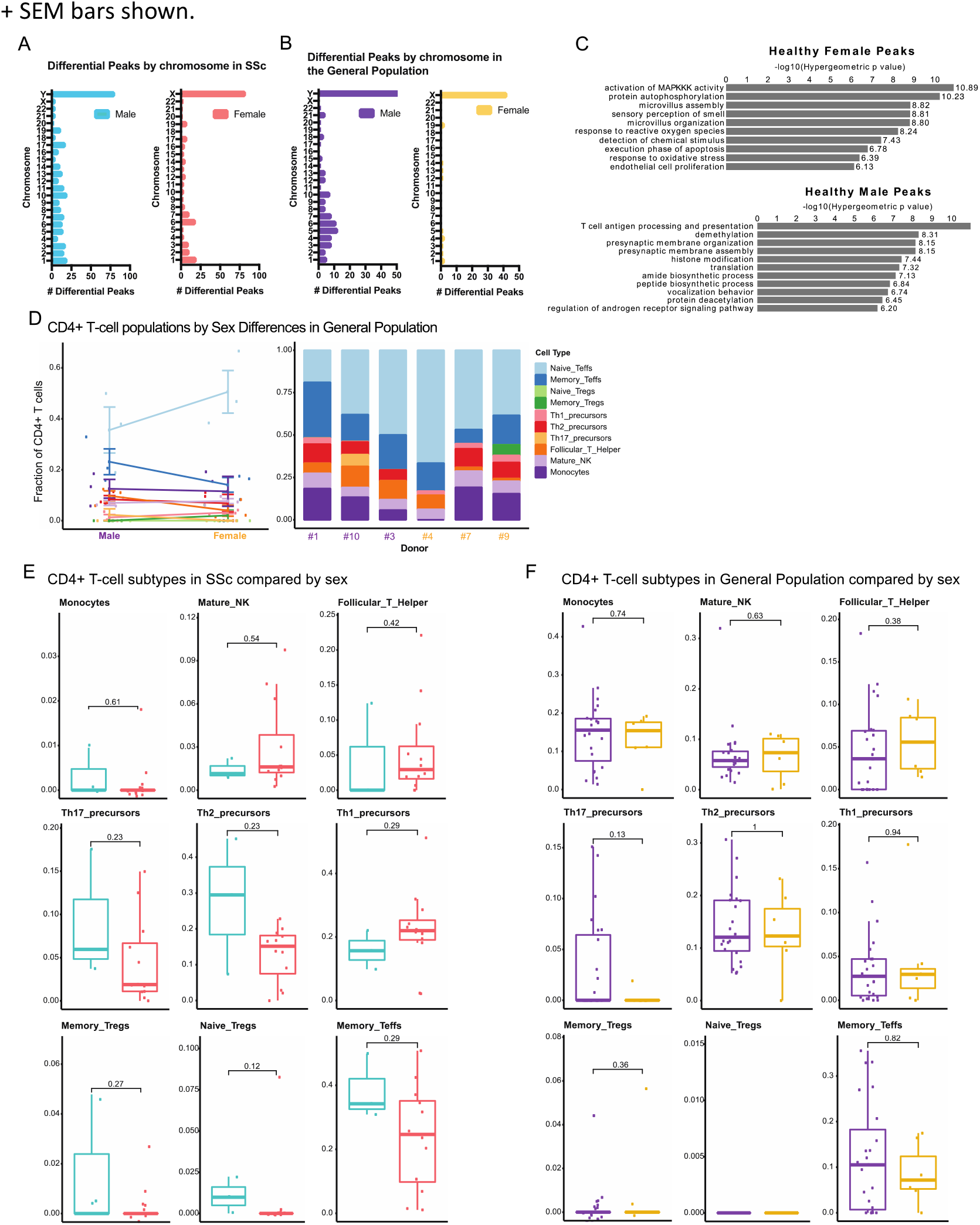
Sex-related differences in SSc study patients. **A** Differential peaks from each chromosome grouped by sex (female = red, male = blue) in SSc study patients. **B** Differential peaks from each chromosome grouped by sex (female = yellow, male = purple) in general population healthy control samples. **C** GREAT predicted functions of cis-regulatory regions associated with peaks accessible in healthy control females and males. Statistics calculated using Fisher’s Exact test, 31 total samples from N=7 healthy males and N=3 healthy females. **D** CIBERSORT analysis of CD4^+^ T-cell subsets in general population control males and females (composite of all donors by cell type, left, individual donor examples, right). Percentages of each CD4^+^ T-cell subset as calculated using CIBERSORT compared by sex in **E** SSc patients (N=15 at the baseline timepoint, SSc-1736, - 1753, and -1759 excluded because they had >2 months of treatment at the 1^st^ study timepoint) and **F** general population controls (N=10 donors, 31 samples total). Graphs display Mean + SEM, significance calculated using the Wilcoxon Rank Sum Test.

**Supplemental Figure 4:**
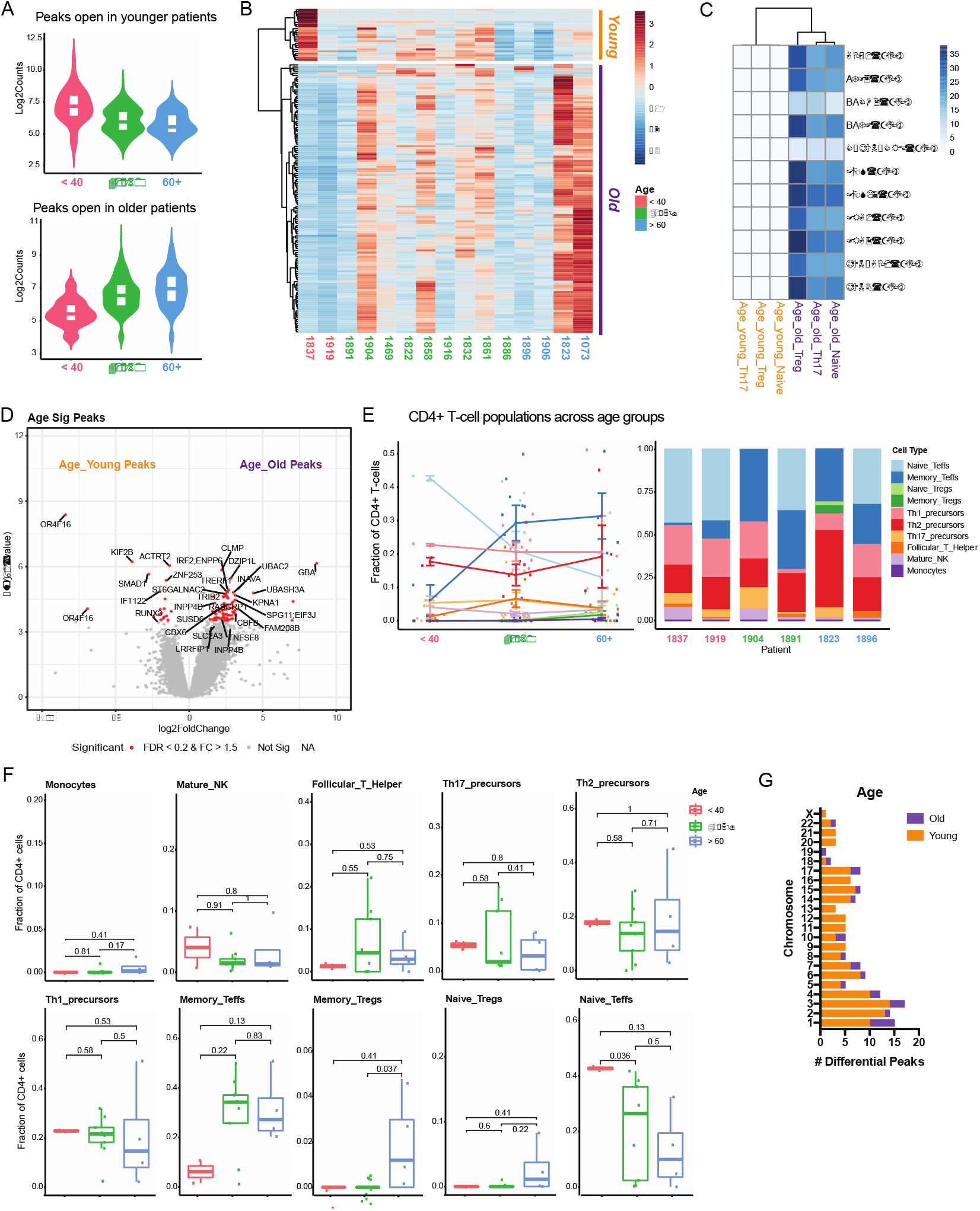
Age-related difference in SSc study patients. **A** Violin plots depicting peaks more accessible in younger than 40 years (red), 41-60 years (green), and older than 60 years (blue) at baseline. **B** Heatmap of z-score for accessibility of differential ATAC-seq peaks of SSc study patients at treatment-naïve, baseline timepoint grouped by age (red label = younger than 40, green label = 41-60 years, blue label = older than 60) and clusters for peaks more accessible in young patients (orange) and old patients (purple). **C** Heatmap of motif enrichment of the genomic region from overlay of differential ATAC-seq peaks with HiChIP anchors. **D** Volcano plot of differential peaks with FDR < 0.2 and absolute fold change larger than 1.5 highlighted in red. “Young” (orange) and “Old” (purple) refer to the two clusters from **B. E** CIBERSORT analysis of CD4^+^ T-cell subsets grouped by age of SSc patients (left panel: composite of N=17 patients by cell type; right panel: individual patient examples). **F** Percentages of each CD4^+^ T-cell subset as calculated by CIBERSORT compared by age group in SSc patients. **G** Differential peaks from each chromosome grouped by age in SSc study patients. Significance calculated using the Wilcoxon Rank Sum Test, N=15 patients at the baseline timepoint. SSc-1736, -1753, and -1759 excluded because they had >2 months of treatment at the 1^st^ study timepoint.

**Supplemental Figure 5:**
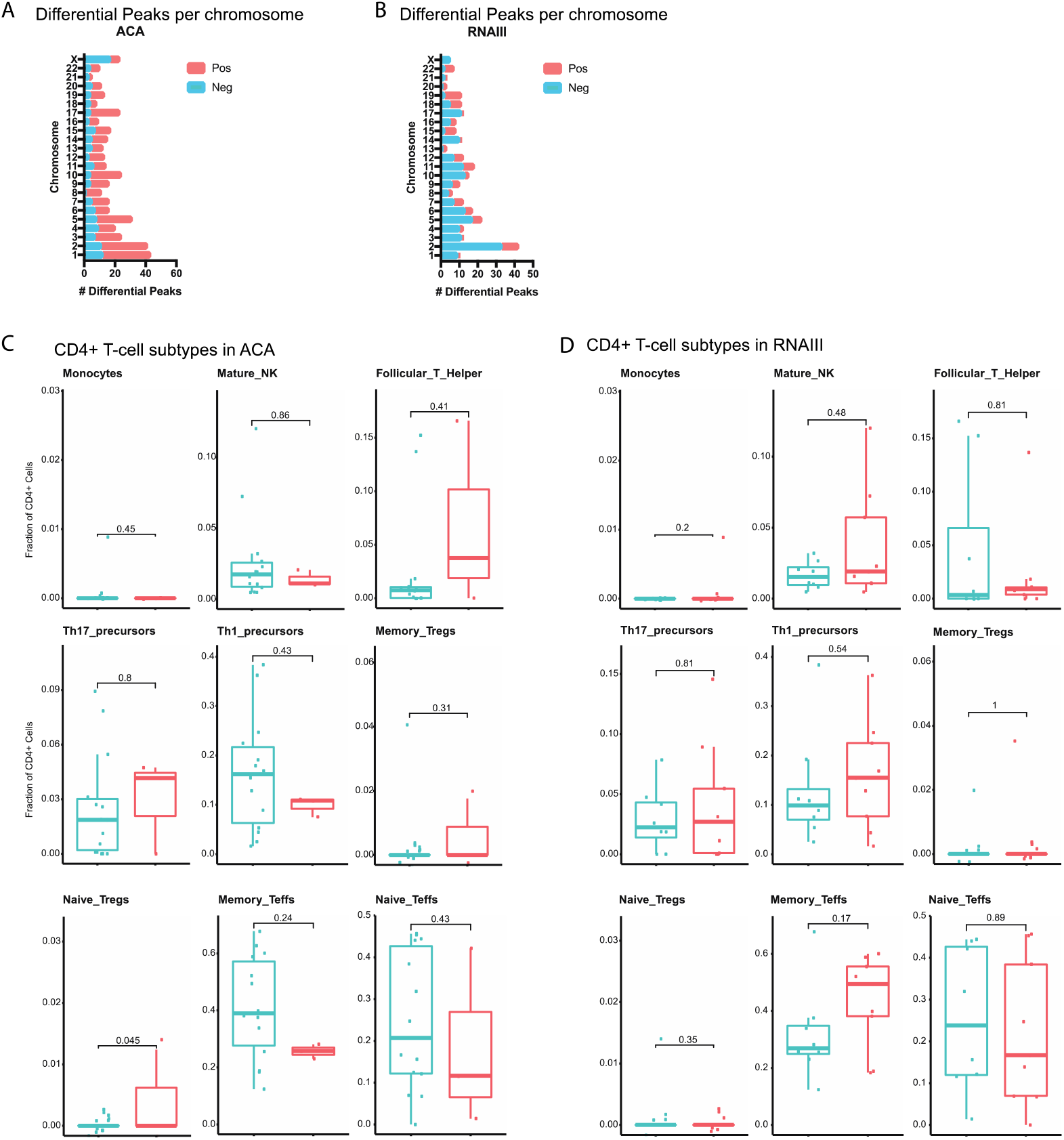
Autoantibody-related differences in SSc study patients. Number of differential peaks at each chromosome grouped by absence and presence of **A** ACA or **B** RNAIII. Percentages of each CD4^+^ T-cell subset as calculated by CIBERSORT compared by positivity or absence of **C** ACA or **D** RNAIII in SSc study patients. Significance calculated using the Wilcoxon Rank Sum Test, N=17 patients at timepoint 3.

